# “Small and many” is the strategy for robust and efficient information transfer in dendritic spines

**DOI:** 10.1101/340414

**Authors:** Takehiro S. Tottori, Masashi Fujii, Shinya Kuroda

## Abstract

A dendritic spine is a small structure on the dendrites of a neuron that processes input timing information from other neurons. Tens of thousands of spines are present on a neuron. Why are spines so small and many? Because of the small number of molecules in the spine volume, biochemical reactions become stochastic. Therefore, we used the stochastic simulation model of N-methyl-D-aspartate receptor (NMDAR)-mediated Ca^2+^ increase to address this issue. NMDAR-mediated Ca^2+^ increase codes the input timing information between prespiking and postspiking. We examined how much the input timing information is encoded by Ca^2+^ increase against prespiking fluctuation. We found that the input timing information encoded in the spine volume (10^-1^ μm^3^) is more robust against prespiking fluctuation than that in the cell volume (10^3^ μm^3^). We further examined the mechanism of the robust information transfer in the spine volume. We demonstrated that the necessary and sufficient condition for robustness is that the stochastic NMDAR-mediated Ca^2+^ increase (intrinsic noise) becomes much larger than the prespiking fluctuation (extrinsic noise). The condition is satisfied in the spine volume, but not in the cell volume. Moreover, we compared the information transfer in many small “spine-volume” spines with that in a single large “cell-volume” spine. We found that many small “spine-volume” spines is much more efficient for information transfer than a single large “cell-volume” spine when prespiking fluctuation is large. Thus, robustness and efficiency are two functional reasons why dendritic spines are so small and many.

**Significance:** A dendritic spine is a small platform for information processing in a neuron, and tens of thousands of spines are present on a neuron. Why are spines so small and many? Here we addressed this issue using stochastic simulation of NMDAR-mediated Ca^2+^ increase in a spine. We demonstrated that smallness of a spine enables the robust information transfer against input fluctuation, and that many small spines are much efficient for information transfer than a single large cell. This is the first demonstration that shows the advantage of the “small and many” of spines in information processing. The “small and many” strategy may be used not only in spines of a neuron, but also in other small and many intracellular organelles.

## Introduction

A spine is a small structure on the dendrites of a neuron, which processes input timing information from other neurons (1). Dendritic spines are characterized by their small volume and large numbers. For example, the volume of a spine on a rat hippocampal CA1 pyramidal cell is ~0.1 μm^3^, which is 10^4^-fold smaller than the cell body (~1000 μm^3^), and 10,000 spines are present on a single cell (2–4). Why are spines so small and many? In this study, we address this issue by using a stochastic simulation model of N-methyl-D-aspartate receptor (NMDAR)-mediated Ca^2+^ increase.

The most representative receptors for spike-input timing detection for synaptic plasticity on the spines of excitatory neurons are NMDARs (5–7). Activation of NMDARs requires the conjunctive inputs of presynaptic spiking (prespiking) and postsynaptic spiking (postspiking) (8, 9) (**Fig. 1A, B**). Prespiking represents glutamate release from the presynaptic terminals (10, 11), and postspiking is a large increase of membrane potential via backpropagating action potentials (12, 13). The prespiking opens NMDARs, and the postspiking removes the receptors’ Mg^2+^ block. Coincident prespiking and postspiking enable the high Ca^2+^ influx via NMDARs (14) (**Fig. 2A, B**). NMDAR-mediated Ca^2+^ increase consequently induces long-term potentiation, which is thought to be the molecular and cellular basis of learning and memory (8, 9, 15).

**Fig. 1.**
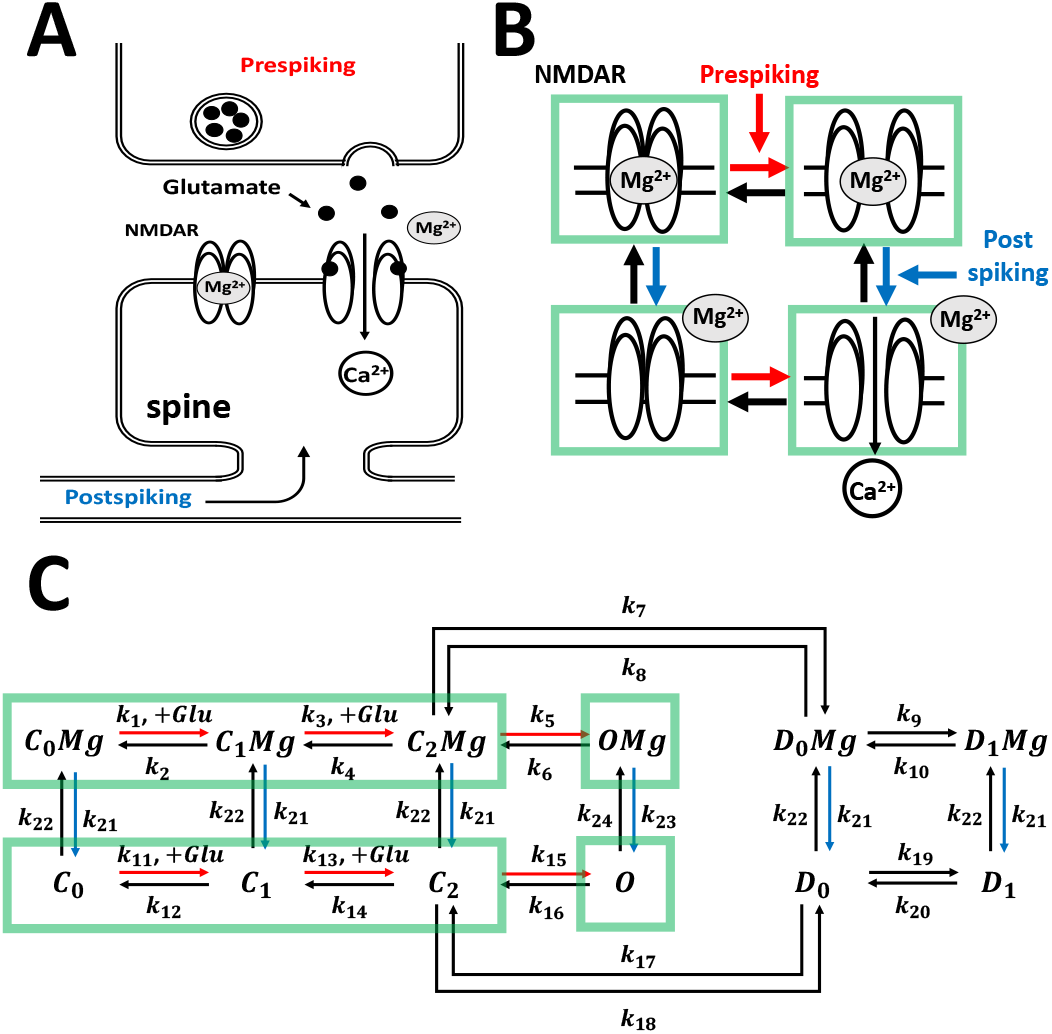
Diagrams of NMDAR-mediated Ca^2+^ increase. (A,B) Schematic diagram of NMDAR-mediated Ca^2+^ increase. Prespiking opens NMDAR, and a postspiking removes the Mg^2+^ block. (C) Block diagram of NMDAR-mediated Ca^2+^ increase. *C*_0_, *C*_1_,*C*_2_,*O*,*D*_0_,*D*_1_ are the states of NMDARs. *C*_0_, *C*_1_, *C*_2_ are the closed states, *O* is the open state, and *D*_0_, *D*_1_ are the desensitization states (see Methods).

The volume of a dendritic spine is so small that the number of molecules in the spine is limited to tens to hundreds. Therefore, biochemical reactions are stochastic in the spine volume, but they are deterministic in the cell volume (16, 17). For example, the number of NMDARs on a spine is estimated to be only about 20 (18–20), which increases the stochasticity in biochemical reactions, and thus the number of NMDARs opening fluctuates widely, which causes large fluctuation in the NMDAR-mediated Ca^2+^ increase (21–23). Intuitively, such a large fluctuation caused by the small volume of the spine is disadvantageous for robust information transfer. Why is the spine so small?

Other representative receptors for spike-timing detection for synaptic plasticity are metabotropic glutamate receptors (mGluRs) (7, 24). We previously demonstrated that mGluR-mediated Ca^2+^ increase is robust against input fluctuation by using a stochastic simulation model of mGluR-mediated Ca^2+^ increase in the cerebellar Purkinje spine (16). The input timing information encoded to Ca^2+^ increase does not decrease much, even when the input fluctuation is large. The robustness appears only in the spine volume rather than the larger volume including the cell volume. mGluR-mediated Ca^2+^ increase is induced by conjunctive inputs of the parallel fiber and climbing fiber (16, 17). Parallel fiber input is glutamate release, leading to activation of mGluR and subsequent production of IP_3_. Climbing fiber input is backpropagating membrane potential, leading to Ca^2+^ increase via voltage-gated Ca^2+^ channels. Only in the presence of IP**3** triggered by parallel fiber input does this Ca^2+^ increase via voltage-gated Ca^2+^ channels trigger the regenerative cycles of Ca^2+^-induced Ca^2+^ release from IP**3** receptors, leading to a large Ca^2+^ increase. In contrast to mGluR-mediated Ca^2+^ increase, which involves multiple biochemical reactions, activation of NMDARs is a single-molecule reaction that depends on glutamate-binding by prespiking and removal of Mg^2+^-block by postspiking. Thus, it remains unclear whether NMDAR-mediated Ca^2+^ increase shows a robustness similar to that of mGluR-mediated Ca^2+^ increase despite the different mechanism. Moreover, the question of why spines are many has not been examined in previous studies (16, 17).

In this study, using a stochastic simulation model of NMDAR-mediated Ca^2+^ increase, we demonstrate that NMDAR-mediated Ca^2+^ increase shows robust information transfer only in the spine volume, but not in the cell volume. The robustness occurs when the extrinsic noise caused by the prespiking fluctuation is much smaller than the intrinsic noise caused by the fluctuation of biochemical reactions. We also compared the information transfer in many small spines with that in a single large spine, and our results demonstrate that many small spines realize efficient information transfer when prespiking fluctuation is large. These results indicate that “small and many” is the strategy for robust and efficient information transfer in dendritic spines.

## Results

### Stochastic simulation model of NMDAR-mediated Ca^2+^ increase

We constructed the stochastic simulation model of NMDAR-mediated Ca^2+^ increase on the basis on the literature (25–32) (**Fig. 1C**). Initial states and reaction constants are described in Tables S1 and S2. In this model, prespiking and postspiking are the inputs, and Ca^2+^ is the output. *C*_0_,*C*_1_,*C*_2_,*O*,*D*_0_,*D*_1_ are the states of NMDARs. *C*_0_,*C*_1_,*C*_2_ are the closed states, *O* is the open state, and *D*_0_, *D*_1_ are the desensitization states. Mg^2+^ can bind all states of NMDARs. The prespiking is glutamate, and postspiking is backpropagating action potential. Prespiking induces the transition from the closed states to the open state, and postspiking induces the dissociation of Mg^2+^ from NMDARs. Based on our previous deterministic model of NMDAR-mediated Ca^2+^ increase (27), we simulated NMDAR-mediated Ca^2+^ increase using a stochastic simulation algorithm (see Methods).

### NMDAR-mediated Ca^2+^ increase in the spine and cell volumes

We examined the time course of Ca^2+^ increase in the spine volume and cell volume using the stochastic simulation model of the NMDAR-mediated Ca^2+^ increase (**Fig. 2**). Hereafter, we denoted 10^-1^ μm^3^ as the spine volume and 10^3^ μm^3^ as the cell volume. When postspiking occurred at 10 msec after prespiking, a large Ca^2+^ increase was always observed in the spine volume and in the cell volume (**Fig. 2C, D**), consistent with the earlier experimental result (14) (**Fig. 2A, B**). However, the time course of Ca^2+^ increase was always the same in the cell volume, whereas it differed greatly between trials in the spine, indicating that Ca^2+^ increase was deterministic in the cell volume and stochastic in the spine volume.

**Fig. 2.**
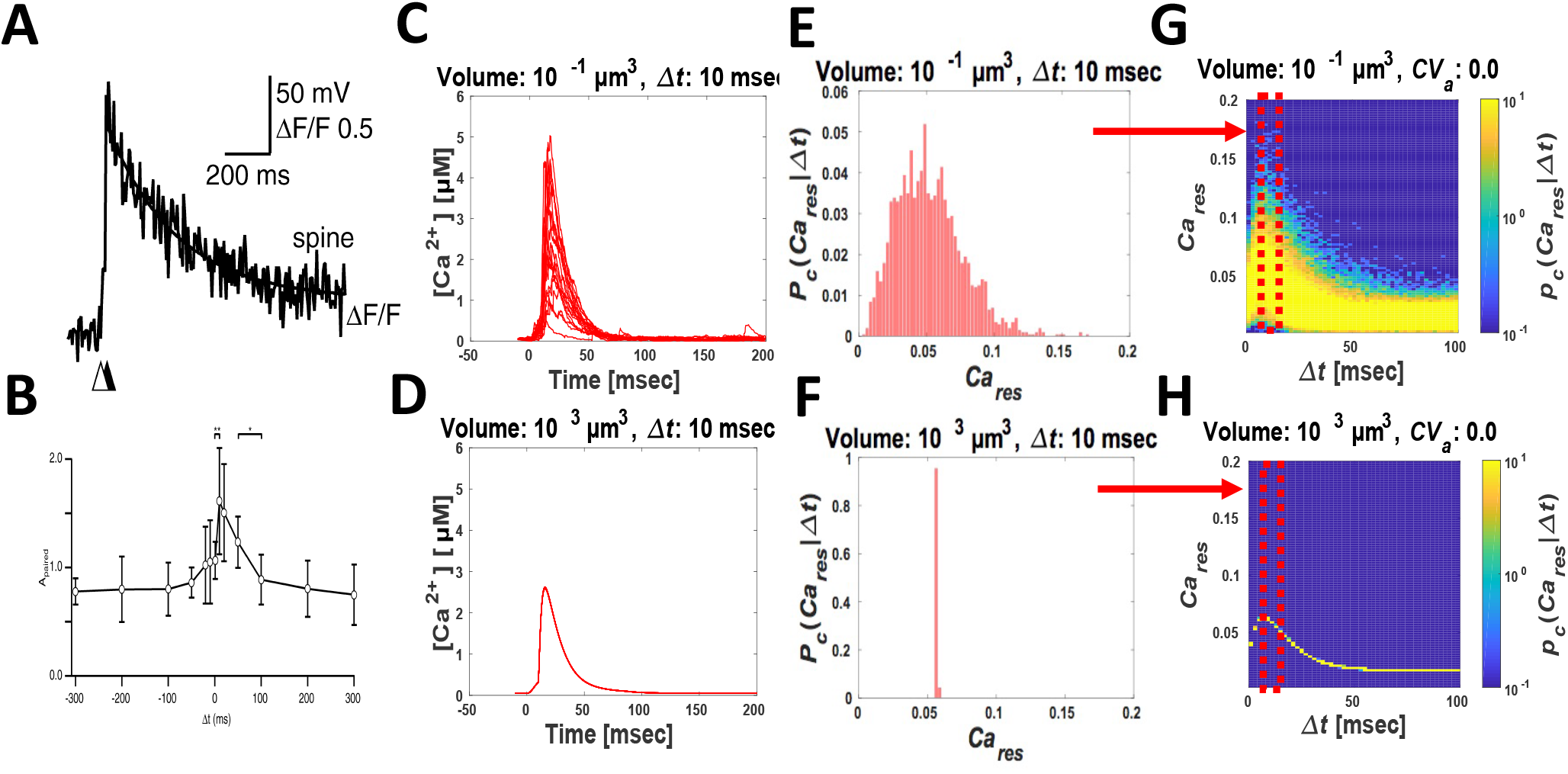
NMDAR-mediated Ca^2+^ increase in the stochastic model. (A,B) Experimental results of NMDAR-mediated Ca^2+^ increase by prespiking and postspiking(14). (A) Time course of Ca^2+^ increase at the pre-postspiking interval *Δt* = *t_post_* − *t_pre_* = 10 msec. (B) Peak of Ca^2+^ increase with the pre-postspiking interval Δt. (C-H) Stochastic simulation results of NMDAR-mediated Ca^2+^ increase in the spine volume (10^-1^ μm^3^) (C,E,G) and in the cell volume (10^3^ μm^3^) (D,F,H). (C,D) Time-courses of Ca^2+^ increase in the spine volume (C) and in the cell volume (D) at the pre-postspiking interval *Δt* = 10 msec (number of trials is 20). (E,F) Probability density distributions of *Ca_res_* with the pre-postspiking interval *Δt*, *p_c_*(*Ca_res_*|*Δt*), in the spine volume (E) and in the cell volume (F) at *Δt* = 10 msec (number of trials is 2000). (G,H) Probability density distributions of *Ca_res_* with the pre-postspiking interval *Δt*, *p_c_*(*Ca_res_*|*Δt*), in the spine volume (G) and in the cell volume (H) from *Δt* = 0 msec to *Δt* = 100 msec (number of trials is 2000).

To quantify the variability of Ca^2+^ increase, we examined the probability density distribution of *Ca_res_*, which was defined by the temporal integration of Ca^2+^ concentration, subtracted by the basal Ca^2+^ concentration. The probability density distribution of *Ca_res_* was unimodal in both the spine and cell volumes (**Fig. 2E, F**). We previously reported that the probability density distribution of mGluR-mediated Ca^2+^ increase was unimodal in the cell volume, whereas it was bimodal in the spine volume (16, 17), indicating that the probability density distribution of *Ca_res_* in the spine volume differs between NMDAR-mediated and mGluR-mediated Ca^2+^ increase.

Next, we examined the probability density distribution of *Ca_res_* at 2-msec pre-postspiking intervals from 0 to 100 msec. The probability density distribution of Ca_res_ was unimodal regardless of pre-postspiking intervals in both the spine and cell volume (**Fig. 2G, H**). In addition, the mean of the probability density distribution of *Ca_res_* changed depending on the pre-postspiking interval. For example, when postspiking occurred within 30 msec after prespiking, the mean of the probability density distribution of *Ca_res_* became large in the spine volume and in the cell volume.

### Robustness against prespiking fluctuation in the spine volume

The amplitude of prespiking, the released glutamate concentration, was different between trials (22, 33). Thus, the fluctuation of Ca^2+^ increase can be influenced by the prespiking fluctuation as well as the stochasticity in biochemical reactions. The former can be regarded as extrinsic noise, and the latter as intrinsic noise. We examined the effect of prespiking fluctuation on the fluctuation of the Ca^2+^ increase.

The distribution of the amplitude of prespiking can be approximated by the Gaussian distribution (34). Therefore, we set the probability density distribution of the amplitude of prespiking *p_a_*(*Amp_pre_*|*μ_a_*, *σ_a_*) as the Gaussian distribution given by *N*(*Amp_pre_*|*μ_a_*,*σ_a_*), where *Amp_pre_* denotes the amplitude of prespiking and *μ_a_* and 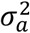 denote the mean and the standard deviation of the amplitude of prespiking in each trial, respectively.

We examined the probability density distribution of *Ca_res_* against the coefficient of variation of the amplitude of prespiking, *CV_a_*. Note that *CV_a_* is *σ_a_*/*μ_a_* with *μ_a_* = 6.0 μM. In the spine volume, the probability density distributions of *Ca_res_* with *CV_a_* = 0.0 (**Fig. 3A**) and *CV_a_* = 0.5 (**Fig. 3B**) were similar; however, as the volume increased, the probability density distributions of *Ca_res_* with *CV_a_* = 0.0 (**Fig. 3C, E**) and *CV_a_* = 0.5 (Fig. 3D, F) changed enormously. The variability of *Ca_res_* remained similar regardless of *CV_a_* in the spine volume, whereas the variability of *Ca_res_* increased as the *CV_a_* increased in the cell volume (**Fig. S1**).

**Fig. 3.**
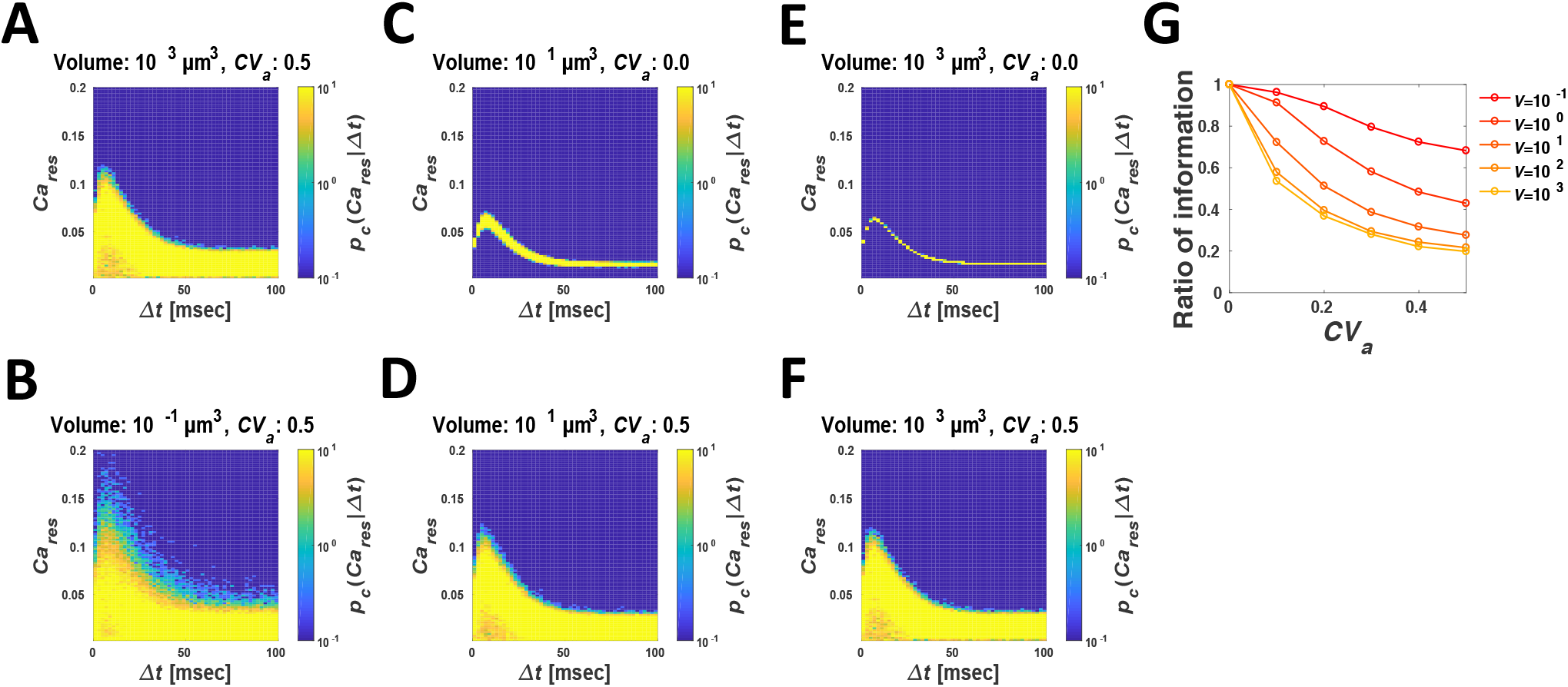
Probability density distributions of *Ca_res_* against the prespiking fluctuation. (A-F) Probability density distributions of *Ca_res_* in the spine volume (A,B), in the intermediate volume (10^1^ μm^3^) (C,D), and in the cell volume (E,F) with the coefficient of variation of prespiking *CV_a_*. *CV_a_* = 0 (A,C,E) and *CV_a_* = 0.5 (B,D,F). (G) The mutual information between the pre-postspiking interval *Δt* and the *Ca_res_* with the prespiking fluctuation *CV_a_* in the indicated volume. The mutual information is normalized by the mutual information at *CV_a_* = 0.0 for each volume.

Next, we calculated the mutual information of the probability density distribution of *Ca_res_* against the pre-postspiking intervals Δ*t*, *I*(*Ca_res_*|*Δt*), as a measure of how much information of the pre-postspiking interval is transferred to the Ca^2+^ increase (Fig. 3G) (see Methods). In the cell volume, the mutual information *I*(*Ca_res_*|*Δt*) decreased greatly as *CV_a_* increased. In the spine volume, however, the mutual information *I*(*Ca_res_*|*Δt*) did not decrease so much as *CV_a_* increased. Thus, information of the pre-postspiking interval is robustly transferred to the Ca^2+^ increase against the prespiking fluctuation in the spine volume but not in the cell volume.

### Unchanging probability density distributions of *Ca_res_* against prespiking fluctuation in the spine volume explains the robustness

The mutual information *I*(*Ca_res_*|*Δt*) decreased greatly as *CV_a_* increased in the cell volume, whereas *I*(*Ca_res_*|*Δt*) did not decrease as much as *CV_a_* increased in the spine volume (**Fig. 3G**). Why was the mutual information in the spine volume more robust against *CV_a_* than that in the cell volume?

The change of the probability density distribution of *Ca_res_* against *CV_a_* in the spine volume was smaller than that in the cell volume (**Fig. 3A–F**). Generally, when an input distribution and an output distribution do not change, the mutual information between the input and output remains the same. In other words, the change of the mutual information *I*(*Ca_res_*|*Δt*) can be caused by the change of the probability density distribution of *Ca_res_* against *CV*_a_.

Therefore, we examined the change of the probability density distribution of *Ca_res_* against *CV_a_* in more detail, calculating the probability density distribution of *Ca_res_* with the fluctuation of *Amp_pre_*, *p_ac_*(*Ca_res_*|*μ_a_, σ_a_*) (**Fig. 4B, D, F**). Note that the pre-postspiking interval *Δt* was fixed. We previously set the probability density distribution of *Amp_pre_, p_a_*(*Amp_pre_*|*μ_a_,σ_a_*) as the Gaussian distribution *N*(*Amp_pre_*|*μ_a_,σ_a_*), based on experimental observations (34). In addition, we obtained the probability density distribution of *Ca_res_* against *Amp_pre_, p_c_*(*Ca_res_*|*Amp_pre_*), from the stochastic simulation (**Fig. 4A, C, E**). Thus, the probability density distribution of *Ca_res_* with the fluctuation of *Amp_pre_, p_ac_*(*Ca_res_*|*μ_a_,σ_a_*) is given by the following:

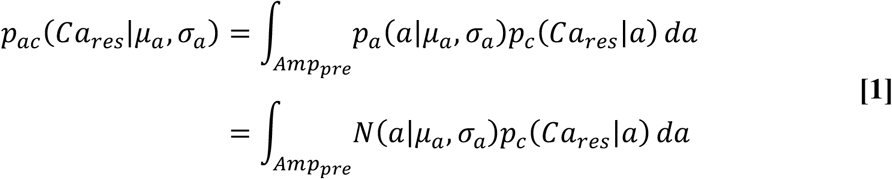

(**Fig. 4B, D, F**). Here, *a* ∈ *Amp_pre_*. In the spine volume, the probability density distribution of *Ca_res_* remained similar with an increase in the *CV_a_* (**Fig. 4B**). In larger volumes including the cell volume, the probability density distribution of *Ca_res_* changed greatly as *CV_a_* increased (**Fig. 4D, F**).

**Fig. 4.**
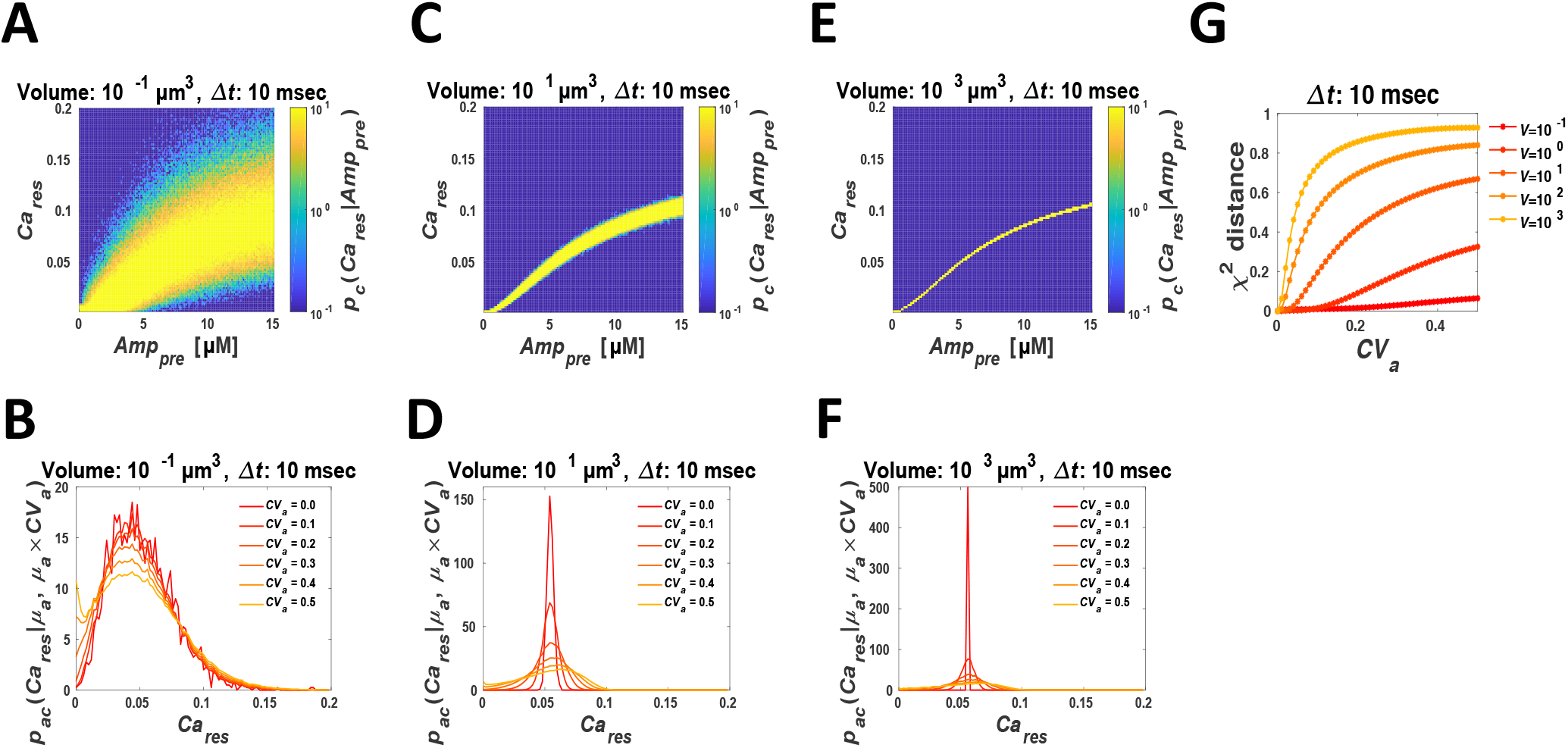
Changes of probability density distributions of *Ca_res_* with the increase in prespiking fluctuation. (A,C,E)**e** Probability density distributions of *Ca_res_* against *Amp_pre_* in the spine volume (A), in the intermediate volume (C), and in the cell volume (E) at the pre-postspiking interval *Δt* = 10 msec. *Amp_pre_* indicates the amplitude of prespiking. (B,D,F) Probability density distributions of *Ca_res_* against the prespiking fluctuation *CV_a_* in the spine volume (B), in the intermediate volume (D), and in the cell volume (F) at the pre-postspiking interval *Δt* = 10 msec. (G) The *CV_a_*-dependency of *χ*^2^ distance between the probability density distribution of *Ca_res_* at *CV_a_* and that of *Ca_res_* at *CV_a_ =* 0.0.

Next, we quantified the change of the probability density distribution of *Ca_res_* against the *CV_a_* based on the *χ*^2^ distance between the probability density distribution of *Ca_res_* at *CV_a_* = 0.0 and the probability density distribution of *Ca_res_* with the increase in the *CV_a_* (**Fig. 4G**). The χ^2^ distance is 0 when two distributions are the same, and it is 1 when two distributions are completely different. In the cell volume, the *χ*^2^ distance increased greatly with the increase in *CV*_a_, and was 0.87 at *CV_a_* = 0.5. In the spine volume, the *χ*^2^ distance remained almost the same regardless of *CV_a_*, and was 0.042 at *CV_a_* = 0.5. Similar results were obtained with other *Δt* such as *Δt* = 30 msec and *Δt* = 70 msec (**Fig. S2**). Thus, the change of the probability density distribution of *Ca_res_* in the spine volume was much smaller than that in the cell volume with an increase in *CV*_a_. This result indicates that unchanging probability density distribution of *Ca_res_* against the prespiking fluctuation in the spine volume, but not in the larger cell volume, is the reason for the robustness

### Larger intrinsic noise than extrinsic noise is critical for the robustness

Why was the probability density distribution of *Ca_res_* against *CV_a_* unchanged in the spine volume, but not in the larger volume? We examined the necessary and sufficient conditions for the robustness, as in the previous study (17). When we define *x* = *Amp_pre_* − *μ_a_*, the relative amplitude of prespiking Eq. 1 changes as follows:

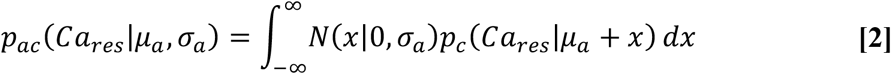

Because *N*(*x*|0,*σ_a_*) is symmetric with respect to *x* = 0, Eq. 2 becomes

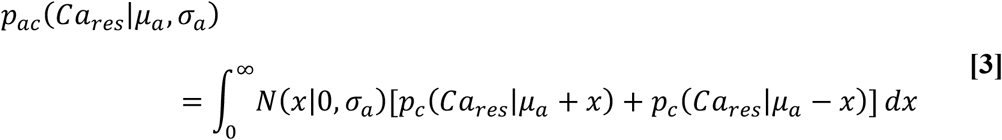

When *x*, the relative amplitude of prespiking, is very large or very small, *N*(*x*|0,*σ_a_*) is also small. In particular, the probability where *x* is included in the range —3*σ_a_* ≤ *x* ≤ 3*σ_a_* is given by 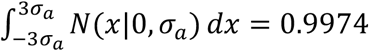, that is, ≈ 1. Thus, the probability of *x* < −3*σ_a_* or *x* > 3*σ_a_* is quite small and almost negligible. Therefore, Eq. 3 can be approximated as follows:

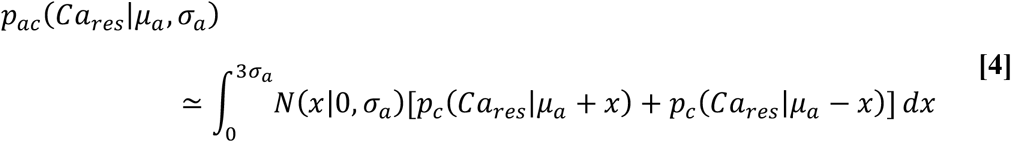

Here, we considered the case given by the following:

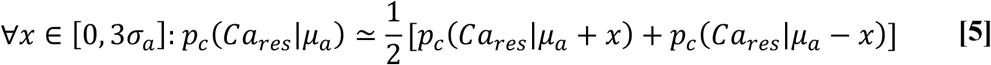

When the conditions where Eq. 5 is satisfied, we obtained the following from Eq. 4:

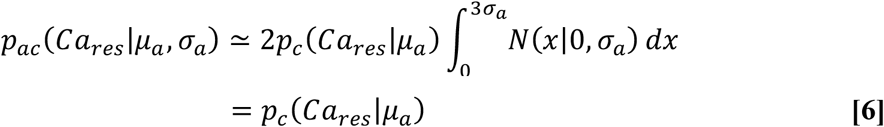

When *CV_a_* = 0.0, that is, *σ_a_* = μ_a_ × *CV_a_* = 0.0, *N*(*a*|*μ_a_,σ_a_*) becomes *δ*(*a*-*μ_a_*), and therefore *p_ac_*(*Ca_res_*|*μ_a_,σ_a_*) becomes *p_c_*(*Ca_res_*|*μ_a_*) at *CV_a_* = 0.0 from Eq. 1. Thus, when Eq. **5** is satisfied, the probability density distribution of *Ca_res_* does not change against *CV*_a_. This indicates that Eq. **5** is the necessary and sufficient conditions for the robustness.

Here, we can also derive the condition where Eq. **5** is satisfied (see Supplemental Note I). The condition where Eq. **5** is satisfied is given by the following:

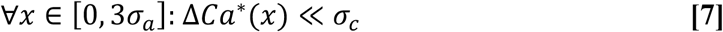

where *ΔCa*\*(*x*) denotes the gap between the peak of *p_c_*(*Ca_res_*|*μ_a_*) and the peak of *p_c_*(*Ca_res_*|*μ_a_* + *x*), which corresponds to extrinsic noise, and *σ_c_* denotes the standard deviation of *p_c_*(*Ca_res_*|*μ_a_*), which corresponds to intrinsic noise. Therefore, the necessary and sufficient condition for the robustness is the range where the extrinsic noise, *ΔCa*\*(*x*), was much smaller than the intrinsic noise, *σ*_c_.

We examined the range of prespiking fluctuation satisfying the condition for robustness. In the spine volume, *ΔCa*\*(*x*)/*σ_c_* ≪ 1 when *x* was small, and *ΔCa*\*(*x*)/ *σ_c_* increased with the increase in *x* (**Fig. 5A**). In contrast, in the cell volume, *ΔCa*\*(*x*)/*σ_c_* exceed 1 even when *x* was small (**Fig. 5A**). Therefore, in the spine volume, information transfer by *Ca_res_* is robust with a larger *CV_a_* because *ΔCa*\*(*x*)/*σ_c_* was smaller than 1 even with a large *x*. In contrast, in the cell volume, information transfer by *Ca_res_* is not robust even with a small *CV_a_* because *ΔCa*\*(*x*)/*σ_c_* was larger than 1 even with a small *x*. We defined *δ_max_* as *x* that gives *ΔCa*\*(*x*)/*σ_c_* = 1. *δ_max_* provides the relative upper bound of *x* where *ΔCa*\*(*x*)/*σ_c_* ≪ 1. Thus, we used *δ_max_* as the index of the range of *x* for robustness (**Fig. 5B**). A larger *δ_max_* means greater robustness. *δ_max_* was highest at the spine volume and decreased with the increase in volume, indicating that the spine volume gives the highest robustness.

**Fig. 5.**
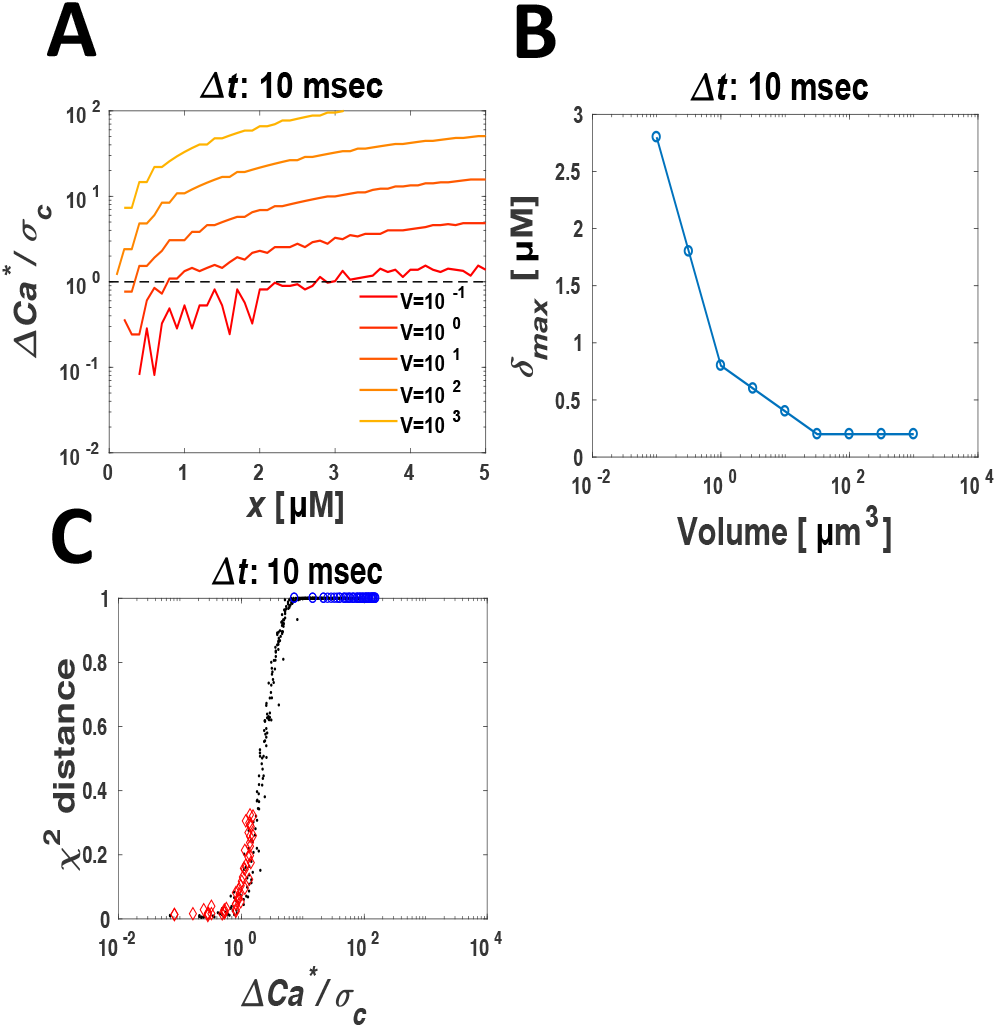
Volume-dependency of the ratio between extrinsic noise and intrinsic noise. (A) The relative amplitude of prespiking, *x*, dependency of *ΔCa*\*/*σ_c_* with *μ_a_ =* 6.0 μM. The dashed line indicates *ΔCa\**/*σ_c_* = 1. The robustness index *δ_max_* is defined as *x* giving *ΔCa\**/*σ_c_* = 1. (B) The volume-dependency of *δ_max_*. (C) The relationship between *ΔCa\**/*σ_c_* and the *χ*^2^ distance between the averaged distribution ^1^ of the probability density distributions of *Ca_res_* with *Amp_pre_* = *μ_a_* ± *x* and that of *Ca_res_* with *Amp_pre_* = *μ_a_*. The red diamonds, blue circles, and black dots indicate the value obtained in the spine volume, cell volume, and intermediate volumes, respectively.

Next, we confirmed that when *ΔCa*\*(*x*)/*σ_c_* is smaller than 1, the probability density distribution of *Ca_res_*, with *Amp_pre_* = *μ_a_, p_c_*(*Ca_res_*|*μ_a_*), and the averaged probability density distribution of *Ca_res_*, with *Amp_pre_* = *μ_a_* ±*x*,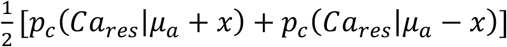, become the same. We quantified the similarities between the two probability density distributions of *Ca_res_* by the *χ*^2^ distance. In the spine volume (**Fig. 5C, *red***), most of the *ΔCa*\*(*x*)/*σ_c_* values were smaller than 1, and the *χ*^2^ distance were also small, indicating that *ΔCa*\*(*x*)/*σ_c_* is smaller than 1 and the two probability density distributions of *Ca_res_* are quite similar in the spine volume. In contrast, in the cell volume (**Fig. 5C, *blue***), most *ΔCa*\*(*x*)/*σ_c_* values were larger than 1 and the *χ*^2^ distances were almost 1, indicating that *ΔCa*\*(*x*)/*σ_c_* is larger than 1, and the two distributions of *Ca_res_* are quite different. Similar results were obtained with other *Δt* such as *Δt* = 30 msec and *Δt* = 70 msec (**Fig. S4**). Therefore, when *ΔCa*\*(*x*), the gap between the two probability density distributions of *Ca_res_*, is smaller than *σ_c_* (i.e., the standard deviation of the probability density distribution of *Ca*_res_), the probability density distribution of *Ca_res_* does not change and becomes robust against fluctuation of prespiking.

### Many small spines versus a single large spine

Why are spines so many? We address this issue with regard to NMDAR-mediated Ca^2+^ increase, which shows more robust information transfer in the smaller volume. By contrast, the mutual information between the pre-postspiking interval and the Ca^2+^ increase decreased as volume decreased (**Fig. 6A**). The mutual information was only 0.45 bit in the spine volume, indicating that the information transfer in the spine is insufficient to reliably determine even a binary decision such as whether prespiking and postspiking are coincident or not. However, spines are many. Around 10,000 spines have been estimated on a single cell (2–4), and all spines are thought to receive the same input, raising the possibility that the entire population of spines code sufficient information of an input.

**Fig. 6.**
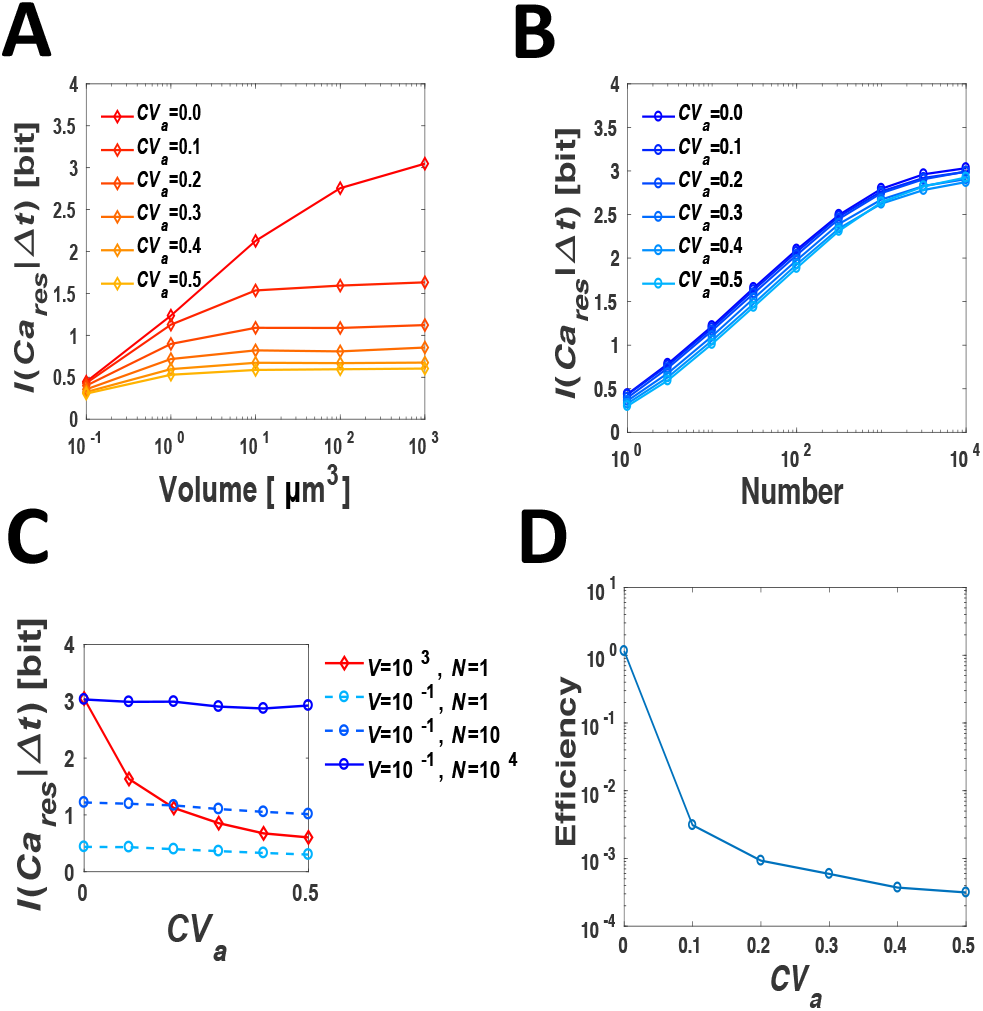
Mutual information in many small spines versus a single large spine. (A) The mutual information between the pre-postspiking interval *Δt* and the *Ca_res_* in I a single large spine. (B) The mutual information between the pre-postspiking interval *Δt* and the *Ca_res_* in many small spines. Their volumes are the spine volume (10^-1^ μm^3^). (C) The *CV_a_*-dependency of the mutual information in a single large “cell-volume” spine (10^3^ μm^3^) (*red*) and in many small “spine-volume” spines (*blue*). (D) Efficiency against *CV_a_*. Efficiency denotes the total volume of many spines (i.e., volume × number) relative to the volume of a cell-volume spine, which can code the: same input timing information.

Therefore, we set the mean of the Ca^2+^ increase over multiple trials as the mean of the Ca^2+^ increase in many spines and examined the mutual information between the pre-postspiking interval and the Ca^2+^ increase in many spines (Fig. 6B). The mutual information increased with the increase in number at *CV_a_* = 0.0. The mutual information was around 1 bit with 3–10 spines at *CV_a_* = 0.0. This result indicates that tens of spines can cooperatively code sufficient input timing information.

We further examined the mutual information of many small spines as *CV_a_* increased. Although the mutual information of a single large spine decreased greatly with the increase in *CV_a_* (**Fig. 6A**), the mutual information of many small spines remained the same regardless of *CV_a_* (**Fig. 6B**). This result indicates that mutual information against the prespiking fluctuation was more robust in many small spines than in a single large spine.

We compared the mutual information against the prespiking fluctuation in many small “spine-volume” spines with that in a single large “cell-volume” spine. In the many small spines, the mutual information remained the same regardless of *CV_a_* (**Fig. 6C *blue***), whereas, in the single large spine, the mutual information abruptly decreased with the increase in *CV_a_* (**Fig. 6C *red***). The mutual information in a cell-volume spine was equal to that in thousands of spines at *CV_a_* = 0.0, while the mutual information in a cell-volume spine was equal to that in only tens of spines at *CV_a_* = 0.5. This result indicates that many small spines are much more efficient for information transfer than a single large cell-volume spine, particularly when prespiking fluctuation is large.

Here, we defined the efficiency as the total volume of many spines (i.e., volume × number) relative to the volume of a cell-volume spine, which can code the same input timing information (**Fig. 6D**). In other words, efficiency means how many spines can code the same input timing information as a cell-volume spine. The efficiency was approximately 1 at *CV_a_* = 0.0 (**Fig. 6D**). This result indicates that, when many small spines code the same input timing information with a cell-volume spine at *CV_a_ =* 0.0, their total volume is equal to the cell volume. However, the efficiency abruptly decreased with the increase of *CV_a_*, and was only 3.1×10^-4^ at *CV_a_* = 0.5 (**Fig. 6D**), indicating that only 3 or 4 spines can code the same amount of input timing information as a cell-volume spine. This result indicates that many small spines are much more efficient for information transfer than a single large cell when the prespiking fluctuation is large.

## Discussion

In this study, we addressed the issue of why dendritic spines are small and many by using a stochastic simulation model of NMDAR-mediated Ca^2+^ increase. We found that smallness of a spine enables robust information transfer of input timing information of pre-postspiking. The robustness appears when the intrinsic noise (i.e., the standard deviation of the Ca^2+^ increase) is larger than the extrinsic noise (the change of the peak of the Ca^2+^ increase caused by the prespiking fluctuation). The intrinsic noise in the spine volume was much larger than that in the cell volume, and thus the information transfer in the spine volume is more robust than that in the cell volume.

We also found that many small spines are much more efficient for information transfer than a single large spine. The input timing information a single small spine can code is much less than 1 bit, but many small spines can cooperatively code sufficient input timing information. In addition, we also demonstrated that many small spines can code the same input timing information as a cell-volume spine, although their total volume is much smaller than the cell volume.

The robust information transfer in the spine volume also can be realized in mGluR-mediated Ca^2+^ increase (Fig. S6) (16, 17), which has a different biochemical mechanism than that of NMDAR-mediated Ca^2+^ increase. This result indicates that the robust information transfer in a small volume is independent of structures of biochemical reactions and a conserved feature in the spines of excitatory neurons.

In addition to the robustness, mGluR-mediated Ca^2+^ increase shows efficiency in information transfer, sensitivity to lower amplitude of input, and probability coding of input timing information (Fig. S6) (16, 17). Note that in the previous study efficiency denotes the mutual information per input molecule efficiency, which is different from efficiency in this study. We also examined whether NMDAR-mediated Ca^2+^ increase shows these properties (see Supplemental Note II). Although NMDAR-mediated Ca^2+^ increase shows efficiency, but it does not show sensitivity and probability coding (Fig. S6). Therefore, efficiency in the previous study is a conserved feature between NMDAR- and mGluR-mediated Ca^2+^ increase, whereas sensibtivity and probability coding are not.

The properties of mGluR such as sensitivity and probability coding that are not seen in NMDAR-mediated Ca^2+^ increase is thought to be caused by the different characteristic of the biochemical reaction: mGluR-mediated Ca^2+^ increase is an excitable system with a positive feedback loop and delayed negative feedback, which shows threshold response, while NMDAR-mediated Ca^2+^ increase is not. We discussed these reasons in more detail in Supplemental Note II. Furthermore, how these different properties between mGluR- and NMDAR-mediated Ca^2+^ increase affect downstream molecules of Ca^2+^ such as CaMKII is not clear, and further investigations are needed.

The robust information transfer in the spine volume can be realized in both NMDAR- and mGluR-mediated Ca^2+^ increase regardless of their different biochemical mechanisms. This indicates that the robustness depends on not biochemical mechanisms but only small volumes. Therefore, robust information transfer may also be seen in other biochemical reactions in spines or in small intracellular organelles, such as intracellular vesicles and mitochondria.

## Methods

### Model

Prespiking represents an increase in glutamate concentration (60 μM), and the glutamate concentration exponentially decreased with the time constant 10 msec (30). Postspiking represents the increase of the membrane potential (60 mV), and the membrane potential exponentially decreased with the time constant 6 msec (31, 32). The resting membrane potential is –65 mV.

In our previous deterministic model of NMDAR-mediated Ca^2+^ increase (27), Ca^2+^ influx via NMDAR was described as follows:

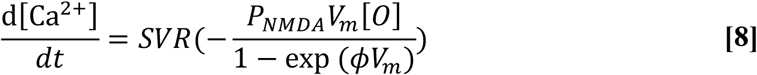

where *SVR* is the surface-to-volume ratio, *V_m_* is the membrane potential, [*O*] is the concentration of NMDAR that is open and free from Mg^2+^, *P_NMDA_* is the permeability of NMDAR channel, and *ϕ* is *2F*/*RT*, where *F* is Faraday’s constant, *R* is the gas constant, and *T* is the absolute temperature (27). Here *SVR* = 11.926 μm^-1^ *P_NMDA_* = 0.05, and *ϕ* = 0.0756 mV^-1^. Furthermore, removal of Ca^2+^ by Ca^2+^ pump was described as follows:

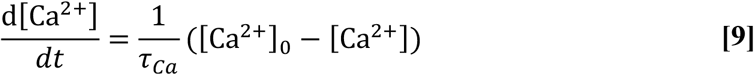

where [Ca^2+^]_0_ is the basal concentration of Ca^2+^, and *τ_Ca_* is the time constant (27). Here *τ_Ca_* = 12 msec (29).

### Stochastic simulation algorithm

These reactions are simulated by the use of the τ-leap method (35). For example, in the reaction represented by Eqs. **8** and **9**, the number of the Ca^2+^ in the spine, *Ca*^2+^(*t* + *τ*), is described as follows:

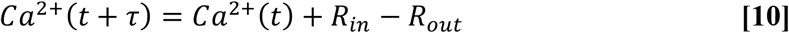

where *R_in_* and *R_out_* indicate the number of reactions of inflow and outflow, which occur in the time interval between *t* and *t* + *τ*, generated to obey the Poisson distribution, 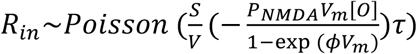 and 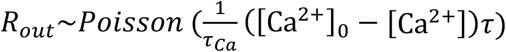, respectively. The appropriate *τ* is calculated in accordance with Cao et al (35).

Under normal circumstances, the reactions by membrane molecules on the membrane, such as NMDAR, and cytosolic molecules, such as Ca^2+^, should be considered as separate mechanisms and compartments, which may be affected by the surface-to-volume ratio. In general, the surface area of the membrane is proportional to the order of the square of length, whereas the cell volume is proportional to the cube of length, which means that, as a system’s size increases, the increasing rate of the number of membrane molecules becomes smaller than that in the cytoplasm. Hence, in the case of larger systems than the spine, the number of membrane molecules that can activate the cytosolic molecules is so small that most substrates are not activated by the stimulation. To identify the simple influences of the smallness of a spine and the number of molecules, it is required that the effect of the stimulation on the cytosolic molecules through the membrane protein for a cell is the same as that for the spine. Therefore, we assumed that the number of membrane proteins is proportional to the volume (i.e., the cube of length) throughout this study.

### Parameter correction

Based on parameters from the literature (25, 26), NMDARs did not bind glutamate much in our model because we assumed that NMDARs are distributed uniformly in the spine to simplify calculation. However, NMDARs are localized to postsynaptic density, and therefore the concentration of NMDARs was underestimated. To calibrate this problem, we increased the reaction constant of NMDAR by 5-fold. However, fewer than 2 NMDARs open, which was smaller than in a previous study (3–5) (21). Therefore, we increased the reaction constant of NMDAR by 5-fold again.

### Mutual information between the input timing and Ca^2+^ increase

We defined *Ca_res_* as the area under the curve of the time course of Ca^2+^, given by the following:

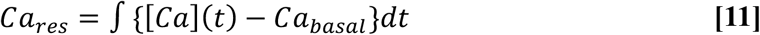

where *Ca_basal_* denotes the basal concentration of Ca^2+^, which is 0.050 μM.

We measured the input timing information coded by the Ca^2+^ increase using the mutual information between the *Ca_res_* and the pre-postspiking interval, *Δt* = *t_post_* − *t_pre_*, given by the following:

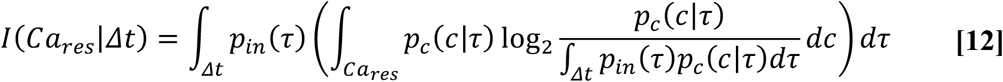

where *p_in_*(*Δt*), the probability density distribution of *Δt*, follows the uniform distribution, and *p_c_*(*Ca_res_*|*Δt*), the conditional probability density distribution of *Ca_res_* given *Δt*, is obtained by the stochastic simulation. Here, *τ* ∈ *Δt* and *c* ∈ *Ca_res_*. To remove the bias caused by the bin width of *Ca_res_*, the mutual information was calculated by using the method introduced by Cheong et al (36).

## Acknowledgements

We thank our laboratory members for their critical reading of this manuscript. This work was supported by The Creation of Fundamental Technologies for Understanding and Control of Biosystem Dynamics, CREST (JPMJCR12W3), from the Japan Science and Technology Agency. M.F. was funded by the Japan Society for the Promotion of Science (JSPS) (JSPS Grants-in-Aid for Scientific Research (KAKENHI) grant No. 6K12508).

## Author contributions

T.S.T., M.F., and S.K. designed research; T.S.T. performed research; T.S.T., M.F., and S.K. analyzed data. T.S.T. and S.K wrote the paper.

## References

1. Greg S, Nelson S, Michael H (2007) Dendrites (Oxford University Press, Oxford, UK).

2. Harris KM, Stevens JK, Harris KM (1989) Dendritic spines of CA1 pyramidal cells in the rat hippocampus: serial electron microscopy with reference to their biophysical characteristics. J Neurosci 9(8):2982–2997.

3. Altemus KL, Lavenex P, Ishizuka N, Amaral DG (2005) Morphological characteristics and electrophysiological properties of CA1 pyramidal neurons in macaque monkeys. Neuroscience. doi:10.1016/j.neuroscience.2005.07.001.

4. Bannister N, Larkman AU (1995) Dendritic morphology of CAl pyramidal neurones from the rat hippocampus: 11. Spine distributions. J Comp Neurol 360:161–171.

5. Malenka RC, Bear MF (2004) LTP and LTD: An embarrassment of riches. Neuron 44:5–21.

6. Lee YS, Silva AJ (2009) The molecular and cellular biology of enhanced cognition. Nat Rev Neurosci 10(2):126–140.

7. Honda M, Urakubo H, Koumura T, Kuroda S (2013) A common framework of signal processing in the induction of cerebellar LTD and cortical STDP. Neural Networks 43:114–124.

8. Bi G, Poo M (2001) Synaptic m odification by c orrelated a ctivity: Hebb’s postulate revisited. Annu Rev Neurosci 24:139–166.

9. Dan Y, Poo M–M (2004) Review spike timing–dependent plasticity of neural circuits. Neuron 44:23–30.

10. Markram H, Tsodyks M (1997) The neural code between neocortical pyramidal neurons depends. Proc Natl Acad Sci U S A 94(January):719–723.

11. Matveev V, Wang XJ (2000) Implications of all–or–none synaptic transmission and short–term depression beyond vesicle depletion: a computational study. J Neurosci 20(4):1575–1588.

12. Hoffman DA, Magee JC, Colbert CM, Johnston D (1997) K+ channel regulation of signal propagation in dendrites of hippocampal pyramidal neurons. Nature 387(6636):869–875.

13. Stuart GJ, Häusser M (2001) Dendritic coincidence detection of EPSPs and action potentials. Nat Neurosci 4(1):63–71.

14. Nevian T (2004) Single spine ca^2+^ signals evoked by coincident epsps and backpropagating action potentials in spiny stellate cells of layer 4 in the juvenile rat somatosensory barrel cortex. J Neurosci 24(7):1689–1699.

15. Sweatt JD (2016) Neural plasticity and behavior - sixty years of conceptual advances. J Neurochem. doi:10.1111/jnc.13580.

16. Koumura T, Urakubo H, Ohashi K, Fujii M, Kuroda S (2014) Stochasticity in Ca^2+^ increase in spines enables robust and sensitive information coding. PLoS One. doi:10.1371/journal.pone.0099040.

17. Fujii M, Ohashi K, Karasawa Y, Hikichi M, Kuroda S (2017) Small–volume effect enables robust, sensitive, and efficient information transfer in the spine. Biophys J. doi:10.1016/j.bpj.2016.12.043.

18. Racca C, Stephenson FA, Streit P, Roberts JD, Somogyi P (2000) NMDA receptor content of synapses in stratum radiatum of the hippocampal CA1 area. J Neurosci 20(7):2512–2522.

19. Sheng M, Hoogenraad CC (2007) The postsynaptic architecture of excitatory synapses: a more quantitative view. Annu Rev Biochem 76:823–847.

20. Okabe S (2007) Molecular anatomy of the postsynaptic density. Mol Cell Neurosci 34(4):503–518.

21. Nimchinsky EA (2004) The number of glutamate receptors opened by synaptic stimulation in single hippocampal spines. J Neurosci. doi:10.1523/JNEUROSCI.5066–03.2004.

22. Hanse E, Gustafsson B (2001) Vesicle release probability and pre–primed pool at glutamatergic synapses in area CA1 of the rat neonatal hippocampus. J Physiol 531(2):481–493.

23. Zeng S, Holmes WR (2010) The effect of noise on CaMKII activation in a dendritic spine during ltp induction. J Neurophysiol 103(4):1798–1808.

24. Ito M (2002) The molecular organization of cerebellar long–term depression. Nat Rev Neurosci 3(11):896–902

25. Kampa BM, Clements J, Jonas P, Stuart GJ (2004) Kinetics of Mg^2+^ unblock of NMDA receptors: implications for spike–timing dependent synaptic plasticity. J Physiol. doi:10.1113/jphysiol.2003.058842.

26. Strecker GJ, Jackson MB, Edward F (1994) Blockade of NMDA–activated channels by magnesium in the immature rat hippocampus. J Neurophysiol 72(4):1538–1548

27. Urakubo H, Honda M, Froemke RC, Kuroda S (2008) Requirement of an allosteric kinetics of NMDA receptors for spike timing–dependent plasticity. J Neurosci. doi:10.1523/JNEUROSCI.0303–08.2008.

28. Sheng M, Hoogenraad CC (2007) The postsynaptic architecture of excitatory synapses: A more quantitative view. Annu Rev Biochem. doi:10.1146/annurev.biochem.76.060805.160029.

29. Sabatini BL, Oertner TG (2002) The life cycle of ca^2+^ ions in dendritic spines. Neuron 33:439–452.

30. Okubo Y, Sekiya H, Namiki S, Sakamoto H, Iinuma S, Yamasaki M, Watanabe M, Hirose K, Iino M (2010) Imaging extrasynaptic glutamate dynamics in the brain. Proc Natl Acad Sci. doi:10.1073/pnas.0913154107.

31. Waters J, Schaefer A, Sakmann B (2005) Backpropagating action potentials in neurones: Measurement, mechanisms and potential functions. Prog Biophys Mol Biol. doi:10.1016/j.pbiomolbio.2004.06.009.

32. Fuenzalida M, Fernández De Sevilla D, Couve A, Buñ W (2010) Role of AMPA and NMDA receptors and back–propagating action potentials in spike timing-dependent plasticity. J Neurophysiol 103:47–54.

33. Branco T, Staras K, Darcy KJ, Goda Y (2008) Local dendritic activity sets release probability at hippocampal synapses. Neuron 59(3):475–485.

34. Isope P, Barbour B (2002) Properties of unitary granule cell-->purkinje cell synapses in adult rat cerebellar slices. J Neurosci 22(22):9668–9678. 35.

35. Cao Y, Gillespie DT, Petzold LR (2006) Efficient step size selection for the tau–leaping simulation method. J Chem Phys. doi:10.1063/1.2159468.

36. Cheong R, Rhee A, Joanne Wang C, Nemenman I, Levchenko A (2011) Information transduction capacity of noisy biochemical signaling networks. Science 334(6054):354–358

